# An inhibitor-interaction intermediate of HIV-1 protease, revealed by Isothermal Titration Calorimetry and NMR spectroscopy

**DOI:** 10.1101/404996

**Authors:** Shahid N Khan, John D Persons, Michel Guerrero, Tatiana V. Ilina, Masayuki Oda, Rieko Ishima

## Abstract

Some of drug-resistant mutants of HIV-1 protease (PR), such as a clinically-relevant drug-resistant PR mutant (Flap+_(I54V)_) containing L10I, G48V, I54V and V82A mutations, produce significant changes in the balance between entropy and enthalpy of the drug-PR interactions, compared to the wild-type (WT) PR. Here, to gain a comprehensive understanding of the entropy-enthalpy compensation effects, we compared nuclear magnetic resonance (NMR), fluorescence spectroscopy and isothermal titration calorimetry (ITC) data of a WT PR with Flap+_(I54V)_ and related mutants: (1) Flap+_(I54V)_; (2) Flap+_(I54A)_ which evolves from Flap+_(I54V)_ in the continued presence of inhibitor yet does not exhibit entropy-enthalpy compensation; and (3) Flap+_(I54)_, a control mutant that contains only L10I, G48V and V82A mutations. Our data indicate that WT and Flap+_(I54A)_ show enthalpy-driven inhibitor-interaction, while Flap+_(I54)_ and Flap+_(I54V)_ exhibit entropy-driven inhibitor interaction. Interestingly, Flap+_(I54A)_ exhibited significantly slower heat flow in the competitive ITC experiment with a strong binder, darunavir, and a weak binder, acetyl-pepstatin, but did not exhibit such slow heat flow in the direct inhibitor-titration experiments. NMR confirmed replacement of the weak binder by the strong binder in a competitive manner. This difference in the heat flow of the competitive binding experiment compared to the direct experiment can only be explained by assuming an inhibitor-bound intermediate pathway. A similar, but attenuated, tendency for slow heat flow was also detected in the competitive experiment with WT. Overall, our data suggests that an inhibitor-bound intermediate affects the entropy-enthalpy compensation of inhibitor-PR interaction.

## Introduction

Human immunodeficiency virus-1 (HIV-1) protease (PR) is an enzyme essential for HIV-1 replication [1-5]. Although structure-based drug design has resulted in the development of various PR inhibitors, the long-term effectiveness of the inhibitors is hampered by generation of drug-resistance mutations [6-20]. To understand the mechanism of the drug-resistance, thermodynamics studies of inhibitor interactions with PR and various drug-resistant mutants have been conducted for the past two decades [21-33]. However, since the PR concentration range typically used in structure studies is much higher than that required for thermodynamics studies, direct comparison of the findings from these two approaches has been hampered. This gap has hindered identification of the protein states responsible for observations made in thermodynamics studies of inhibitor interactions. As a result, modeling of the mechanism of inhibition has been limited.

Our particular interest is Flap+_(I54V)_, which contains a set of clinically-relevant drug-resistant PR mutations, L10I, G48V, I54V and V82A (**Fig. 1**). Thermodynamically, inhibitor interactions, such as saquinavir, amprenavir and darunavir (DRV), with Flap+_(I54V)_ are known to exhibit entropy-enthalpy compensation compared to WT PR: previous isothermal titration calorimetry (ITC) experiments demonstrated that inhibitor-WT interaction was enthalpy driven while inhibitor-Flap+_(I54V)_ was entropy driven [31]. Thus, Flap+_(I54V)_ mutations can silently affect the thermodynamics of drug-PR interactions by changing the entropy and enthalpy balance while not significantly changing the total free energy itself. Interestingly, under drug pressures, Flap+_(I54V)_ evolves to Flap^+^_(I54A)_ (L10I, G48V, I54A and V82A), but DRV interaction with Flap+_(I54A)_ does not exhibit the entropy-enthalpy compensation observed for Flap+_(I54V)_ [32]. Furthermore, the individual mutations, I54V, I54A or G48V, do not show entropy-enthalpy compensation [32]. These observations have indicated that simple addition of each individual mutation effect cannot explain the thermodynamic changes observed for Flap+ mutants [32]. Such cooperativity of the mutations on the inhibitor-binding thermodynamics was also reported for other, similar PR drug-resistance mutants [34].

**Figure 1.**
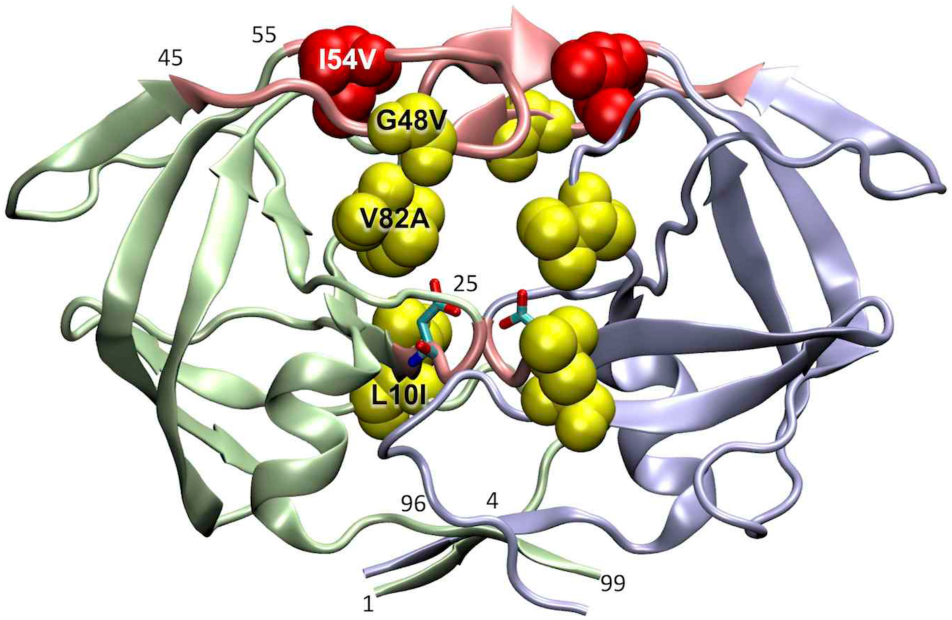
PR structure, showing location of residues L10, G48 and V82 (yellow spheres), and I54 (red spheres) that are mutated in Flap+_(I54V)_ and Flap+_(I54A)_. Two subunits, A and B, are depicted in green and light blue, respectively. Flap region (residues 45-55) and the active-site fireman’s grip (residues 25 to 27) are highlighted in pink with the residue numbers in a small font. Terminal β-sheet regions, residues from 1 to 4 and 96 to 99, are marked in a small font. Structure was generated using PDB: 1T3R [65].

The structure of HIV-1 PR has been well characterized (**Fig. 1**): it is a dimer, with the two subunits interfacing via residues in the flaps (residue 45 - 55), the active-site fireman’s grip (residues 25 to 27) and the N- and C-terminal β-sheet [35-37]. They also indirectly interact with each other through inhibitors at the P1 loop, that include one of the Flap mutation sites, residue 82 (**Fig. 1**). The flaps of HIV-1 PR are known to undergo multiple conformations [38-45]. Previous crystallography, NMR, and MD simulations have shown that Flap+_(I54V)_ exhibits significant fluctuation in the flap region (>0.8 Å Cα rmsd), with more opened flaps in the Flap+_(I54V)_ compared to WT [31; 46; 47], suggesting that two possible apo-forms, i.e., open and closed forms, may contribute to the inhibitor-interaction. Computational and experimental studies have proposed the existence of a PR folding intermediate [48], intermediate-inhibitor bound forms [49], and dimer dissociation as a mechanism of inhibition by DRV [50; 51], which may involve changes of other dimer interface structures. Despite the evidence for multiple conformational states being involved in PR-inhibitor interaction, the thermodynamics of inhibitor- PR interactions have been analyzed assuming only free and bound states.

Recent development in NMR sensitivity have permitted NMR spectra to be recorded for protein samples at low protein concentrations [52], similar to those used for ITC and fluorescence experiments. Taking advantage of this development, we report our ITC, NMR and fluorescence spectroscopy data of PRs, which were recorded at similar conditions, to understand the molecular mechanism underlying PR-inhibitor interaction. We also introduce an artificial mutant, Flap+_(I54)_, which contains mutations L10I, G48V and V82A, and revisit the thermodynamics of inhibitor interaction with Flap+_(I54)_, Flap+_(I54V)_ and Flap+_(I54A)_, as well as that with a pseudo WT (pWT)^1^ protein that is a backbone of the Flap+ mutants (see details in the Materials and Methods), to identify the underlying mechanism of the entropy-enthalpy compensation. We present (i) the thermodynamics parameters for a weak substrate-analogue inhibitor, acetyl-pepstatin (pepstatin), (ii) the apparent thermodynamics parameters of a strong binder, DRV, in the presence of the weak binder, and (iii) the derived thermodynamics parameters of DRV binding. Both pepstatin and DRV, as a weak and a strong binder respectively, have been used to characterize binding thermodynamics of inhibitor-PR interaction in previous studies [22; 26; 27; 31; 32; 53; 54]. To identify the states involved in the PR-inhibitor interaction thermodynamics, we compare Nuclear Magnetic Resonance (NMR) spectra and Trp intrinsic fluorescence emission, at PR concentrations similar to or below those used for ITC. In the end, based on these seamless experiments of both NMR and ITC, we propose that an additional intermediate, possibly an inhibitor-bound dimer intermediate, is needed to explain the data.

## Results

### Pepstatin interaction with PRs monitored by ITC

Pepstatin is an aspartic protease inhibitor and has been used to characterize inhibitor-PR interactions [22; 26; 27; 31; 32; 53-55]. We first characterized the thermodynamics parameters of PR interactions with a weak binder, pepstatin, to utilize the parameters for competitive ITC experiments below and to elucidate inhibitor-interaction characteristics of Flap+ mutants against WT. ITC data of pepstatin with pWT and Flap+ mutants were recorded using similar protein concentrations, 20 – 30 µM. Isotherms of pWT and Flap+_(I54A)_ were similar to each other while isotherms of Flap+_(I54)_ and Flap+_(I54V)_ were similar to each other (**Fig. 2a-h**). Thermodynamics parameters of pepstatin binding to pWT at 20 °C (ΔG, -8.72 ± 0.14 kcal/mol; ΔH, 8.7 ± 0.24 kcal/mol;-TΔS, -17.4 ± 0.28 kcal/mol) were consistent with those obtained at 25 °C by Freire’s group (Δ*G*, -8.4 ± 0.9 kcal/mol; Δ*H*, 10.1± 0.7 kcal/mol, *-T*Δ*S*, -18.4 ± 0.06 kcal/mol) [21] (**Fig.2a** and **2e**, and **Table 1.1**). Thermodynamics parameters of Flap+_(I54A)_ were similar to those of pWT while Flap+_(I54)_ and Flap+_(I54V)_ exhibited less favorable Δ*G* and more unfavorable Δ*H*, compared to pWT and Flap+_(I54A)_ (**Fig. 3a-c** and **Table 1.1**).

**Table 1.**
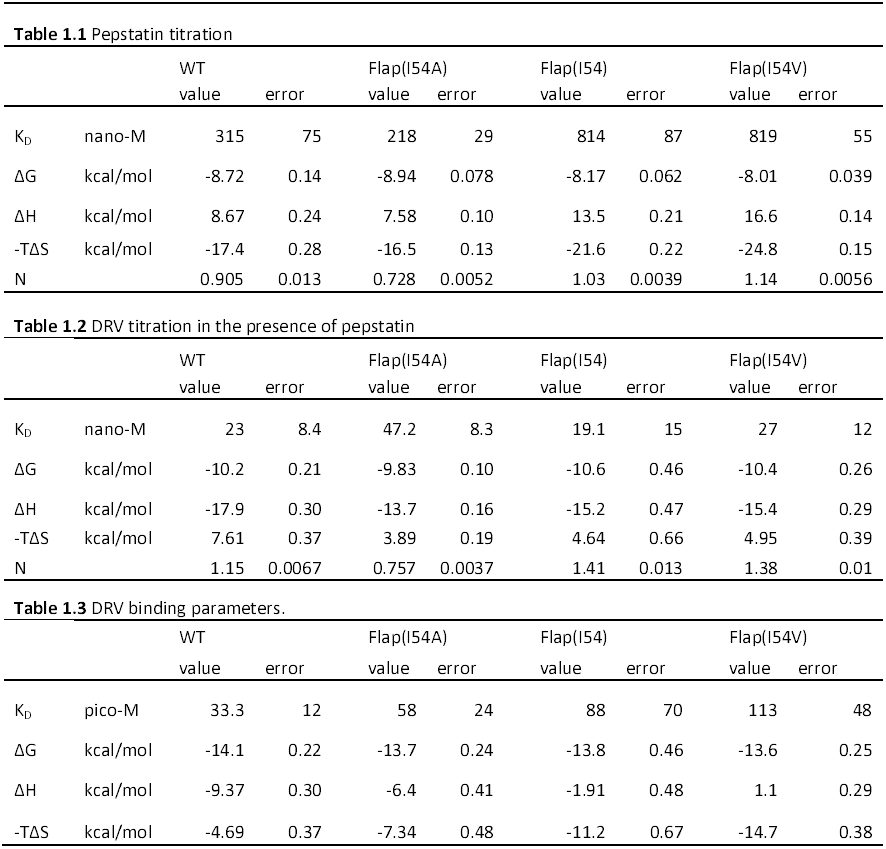
Thermodynamics parameters obtained from calorimetric titration of PR with inhibitors: (1.1) pepstatin-binding parameters, (1.2) DRV-binding parameters in the presence of excess amount of pepstatin, and (1.3) DRV-binding parameters extracted from (1.1) and (1.2).

**Figure 2.**
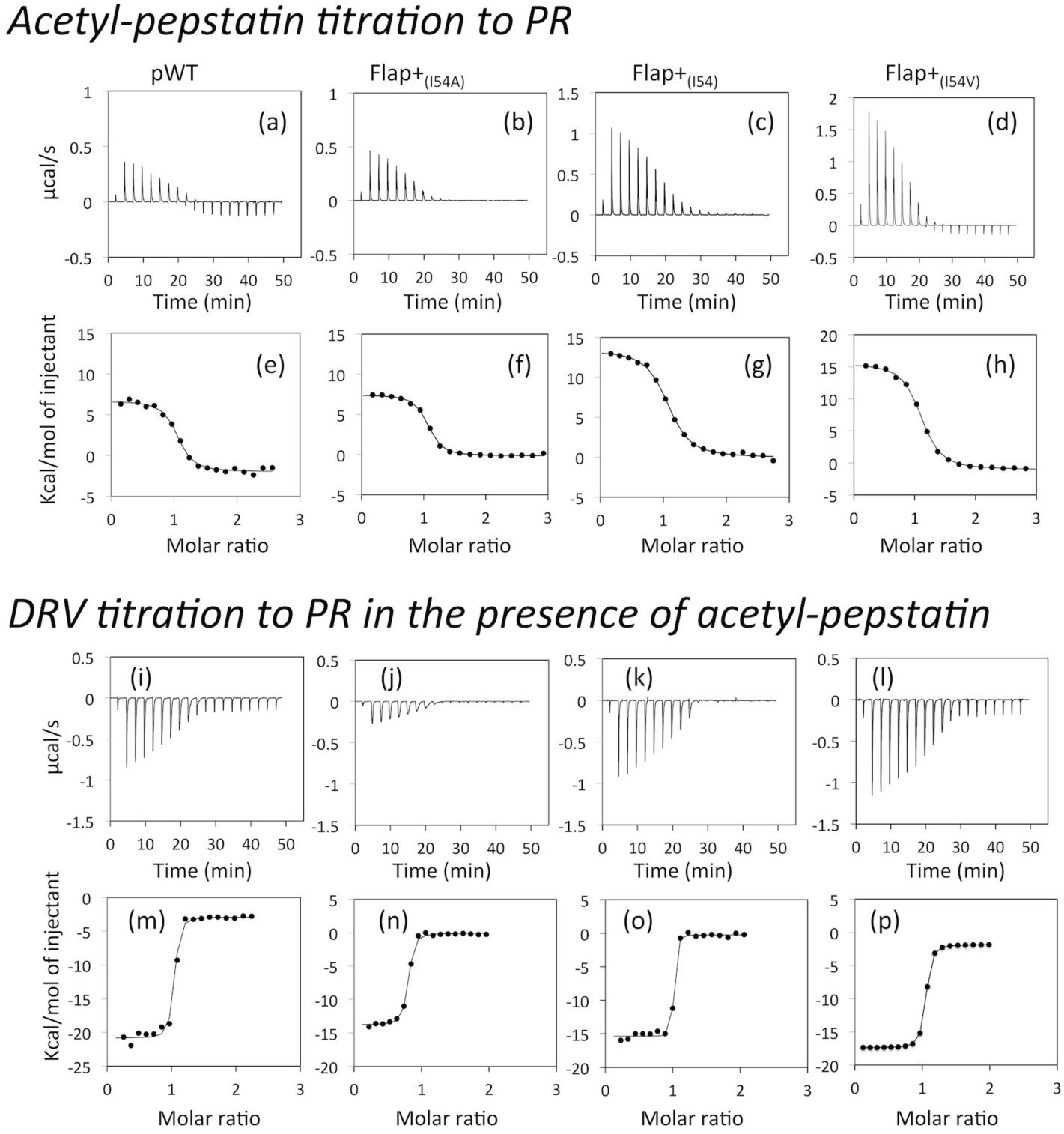
Calorimetric titration of HIV-1 pWT and the Flap+ mutants with (a- h) pepstatin or (i – p) DRV in the presence of excess amount of pepstatin. The heat effects associated with the injection of the inhibitors (a-d and i-l) and corresponding isotherms (e-h and m-p) are shown.

**Figure 3.**
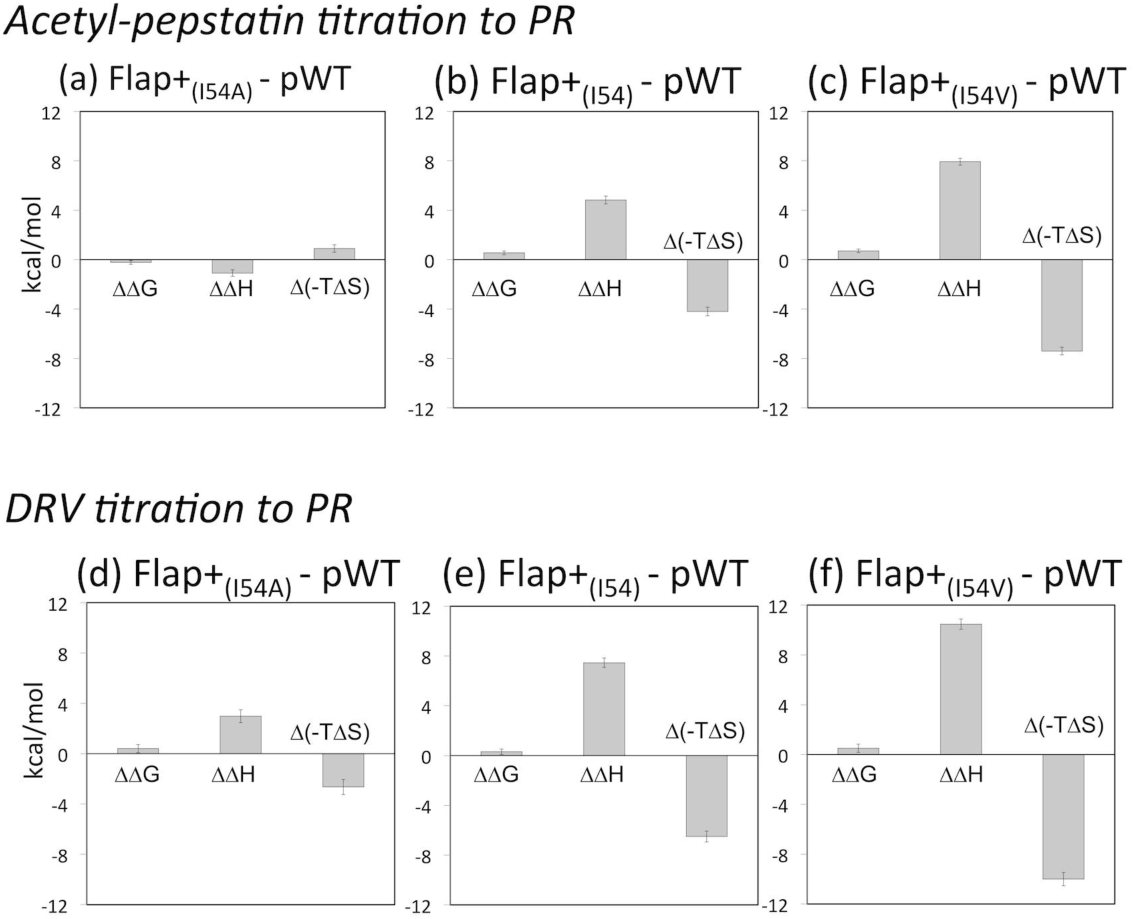
Differences in thermodynamics parameters of Flap+ mutants relative to pWT: (a-c) a- pepstatin binding to the proteins and (d-f) DRV binding to the proteins: (a and d) Flap+_(I54A)_ – pWT; (b and e) Flap+_(I54)_ – pWT; (c and f) Flap+_(I54V)_ – pWT. In each panel, ΔΔ*G* =Δ*G*_Flap+_ - Δ*G*_WT_, ΔΔ*H* =Δ*H*_Flap+_ - Δ*H*_pWT_ and Δ(-*T*Δ*S*) = (-*T*Δ*S*_Flap+_) - (-*T*Δ*S*_pWT_) are shown.

### DRV interaction with PRs monitored by ITC

Titration of PRs with a strong binder, DRV, was done in the presence of a weak binder, pepstatin (**Fig. 2i-2p** and **Table 1.2**). Ideally, the optimal weak binder for competitive ITC experiments would be chosen from a panel of weak binders based on the K_D_ of the strong binder under study [56]. We used a single weak binder, pepstatin, for all PR-DRV interaction studies, to avoid any biases caused by differences in the solubility of weak inhibitors in aqueous solution or differences in the mechanisms of PR interactions of the weak inhibitors. As a result, throughout all DRV-PR interactions, the accuracy of the determined ΔG and *–T*Δ*S* may be less for DRV that shows a steep apparent titration profile. However, Δ*H* is still accurately determined based on the theoretical equation [56]. In the competitive experiments, all the DRV titrations in the presence of pepstatin showed apparent favorable Δ*H* changes, which is in contrast to the pepstatin titration (**Fig. 2i-2p** and **Table 1.2**) but consistent with previous results [32]. As expected, the molar heat changes were steep in the DRV titration for pWT (**Fig. 2m**), indicative that practical Δ*G* accuracy is lower for these proteins.

Using Δ*H* and Δ*G* for the pepstatin titration alone and those of the competitive data (**Table 1.1** and **1.2**), we calculated the thermodynamics parameters of DRV binding to PRs (**Table 1.3**). Favorable Δ*H* was obtained for pWT and Flap+_(I54A)_ while near unfavorable (near zero or positive) Δ*H* values were obtained for Flap+_(I54)_ and Flap+_(I54V)_, respectively (**Table 1.3**). As a result, DRV binding of pWT and Flap+_(I54A)_ was enthalpy-driven while that of Flap+_(I54)_ and Flap+_(I54V)_ was entropy-driven (**Fig. 3d-3f**). The observed Δ*G*, -14.1 ± 0.22 (kcal/mol), of pWT at pH 5.8, which is a condition that slows down autoproteolysis and suitable for a long-term experiments, is slightly higher than that published previously, -15.0 ± 0.3 (kcal/mol), at pH 5.0 [32]. This may be primarily due to inaccuracy of our experimental condition to detect such low Δ*G* binding or secondary because of higher pH in which Asp hydroxyl protonation at the active site may affect to the DRV interaction. Otherwise, thermodynamics parameters of Flap+_(I54A)_ and Flap+_(I54V)_ were similar to those published previously [32].

To reveal the mechanism underlying DRV-pepstatin competition in our experiments, the heat flow of each injection was compared. Since the relative bound fractions at each time point differs among the proteins, both the 3^rd^ and 5^th^ injections were assessed (which corresponds to the ∼7 min and ∼12 min points in **Fig. 4a** and **4b**, respectively). Interestingly, Flap+_(I54A)_ heat flow was quite different from the others. pWT also shows a slightly slower heat flow compared to Flap+_(I54)_ and Flap+_(I54V)_. Importantly, such slow heat flow was not observed (**Fig. 4a** and **4b**, green dashed line) in the direct DRV (**Fig. 4c**) or pepstatin (**Fig. 2b**) titration of Flap+_(I54A)_. Taken together, the thermodynamics changes of inhibitor binding to Flap+_(I54A)_ and pWT are similar to each other, while those of Flap+_(I54)_ and Flap+_(I54V)_ are similar to each other. In Flap+_(I54A)_ andpWT, the competitive experiments are not simply explained by an on and off model of strong and weak binders to PR.

**Figure 4.**
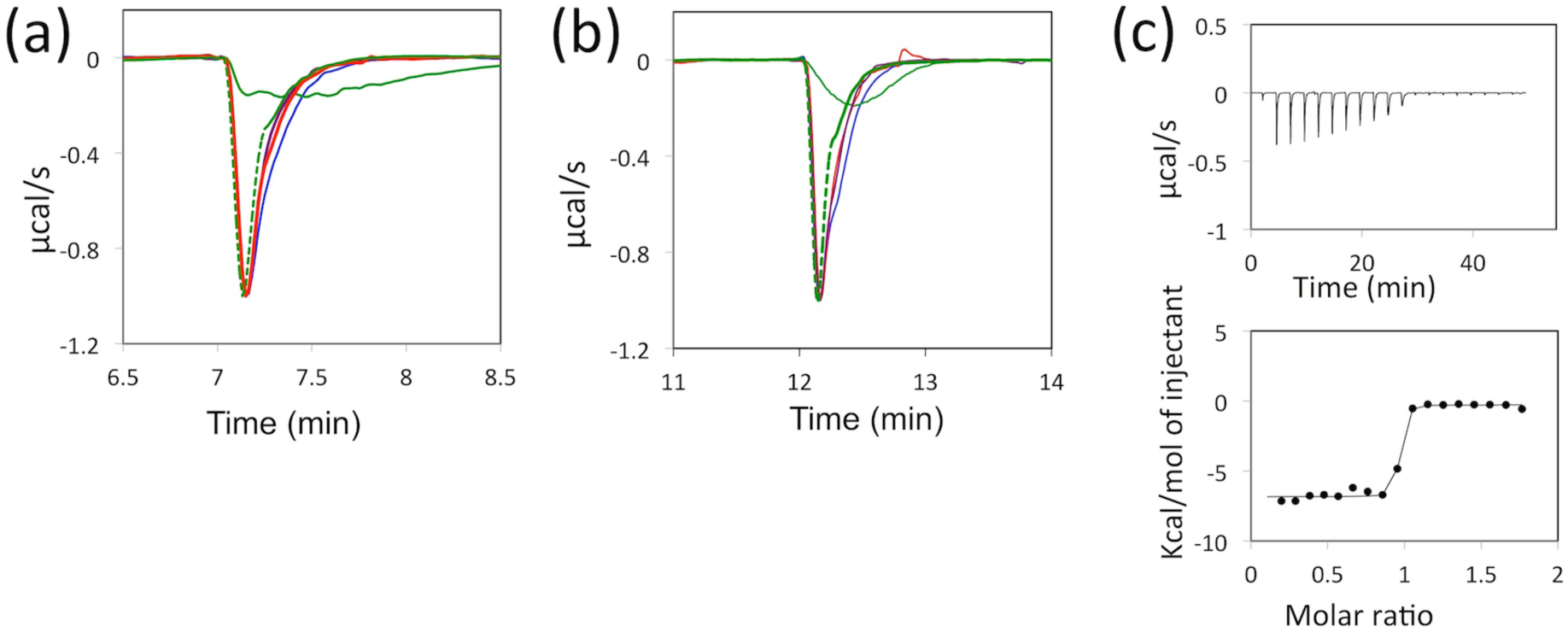
Comparison of heat flow signals of pWT and Flap+ mutants at (a) 7 min and (b) 12 min (3^rd^ and 5^th^ injections) taken from Figures 2i – 2l, and (c) calorimetric titration of Flap+_(I54A)_ with DRV in the absence of pepstatin. In (a) and (b), heat flows of the competitive titration are shown by solid lines for pWT(blue), Flap+_(I54A)_ (green), Flap+_(I54)_ (red), and Flap+_(I54V)_ (purple), while that of the direct Flap+_(I54A)_ titration with DRV is shown by green dashed line. In (a) and (b), maximum heat change was normalized to 1.0 µcal/s to compare the heat flow changes, except for the competitive titration of Flap+_(I54A)_.

### PR dimer dissociation and folding stability

To understand what states are involved in these PR interactions in solution, we next characterized folding and dimerization of the proteins using intrinsic Trp fluorescence spectroscopy at different protein concentrations and denaturant. Since one of the two Trp residues in PR, residue 6, is located at the dimer interface and exposed to solution upon dimer dissociation [57; 58], the fluorescence emission of this residue is known to be reduced by dimer dissociation [48]. In pWT and Flap+ mutants, intrinsic Trp fluorescence spectroscopy exhibited linear protein concentration dependence above 0.25 µM (**Fig. 5a**). Through this analysis, dimer dissociation constants of both pWT and Flap+ mutants were determined to be < 0.25 µM, which is consistent with previous results obtained for PR and other mutants [59] and confirms that the PRs used in our ITC experiments were dimers. Since the reduction of fluorescence emission is lower in Flap+_(I54A)_ compared to pWT, Flap+_(I54)_ and Flap+_(I54V)_, the N-terminal region of Flap+_(I54A)_, where Trp 6 is located, may be more mobile or experience greater monomer unfolding compared to the other proteins.

**Figure 5.**
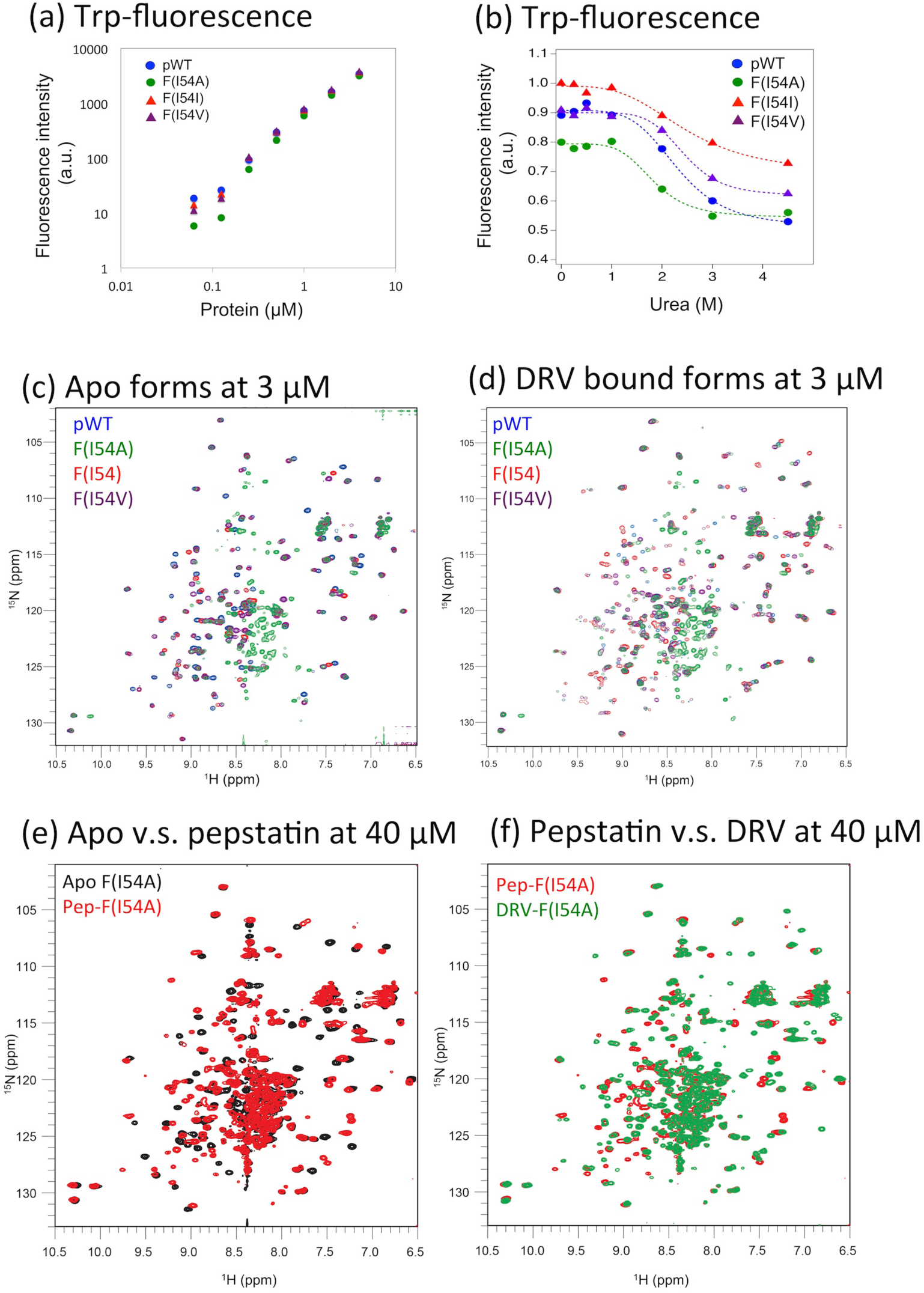
Intrinsic Trp fluorescence emission at 350 nm (a) at varying PR concentrations and (b) at varying urea concentrations, (c, d) overlay of ^1^H-^15^N HSQC spectra of pWT (blue), Flap+_(I54A)_ (green), Flap+_(I54)_ (red), and Flap+_(I54V)_ (purple), in (c) the absence and (d) the presence of DRV at 3 µM protein concentrations, and (e, f) overlay of ^1^H-^15^N HSQC spectra of Flap+_(I54A)_ recorded using a single 40 µM concentration sample (e) before (black) and after (red) adding excess amount of pepstatin, and, (f) in the presence of pepstain, before (red) and after (green) adding excess amount of DRV.

Consistent with this notion, urea denaturation produced overall similar profiles among pWT and Flap mutants (**Fig. 5b**). A slightly lower chemical stability was observed only for Flap+_(I54A)_, with a half urea denaturation concentration at 1.8 ± 0.1 M, compared to the others (2.3 – 2.5 M, respectively) (**Fig. 5b**). In addition, the fluorescence intensity of Flap+_(I54A)_ in the absence of urea was 10 – 20% lower than that of the others, suggesting the presence of an unfolded component or a slightly different conformation of Flap+_(I54A)_ compared to the other PR proteins.

### NMR probed PR conformation

PR structural changes that are involved in the inhibitor-PR interaction observed by ITC were also characterized by recording NMR spectra at different protein concentrations and in the presence and absence of inhibitors. We firstly recorded NMR spectra at a protein concentration lower than those used for ITC, 3 µM, to establish proper folding of the PRs. NMR spectral patterns of pWT and Flap+_(I54V)_ were essentially similar to those published previously, both in apo (**Fig. 5c**) and DRV-bound forms (**Fig. 5d**) [60]. Flap+_(I54)_ spectra also exhibited very similar resonance patterns to those of Flap+_(I54V)_ in both forms (**Fig. 5c** and **5d**). Only the NMR spectrum of Flap+_(I54A)_ showed a significant unfolded fraction, 29% and 17% in the apo and DRV-bound forms, respectively, estimated from the signal intensity of indole NH resonances (**Fig. 5c** and **5d**). These observations of the apo-dimer forms of pWT and Flap+ mutants are consistent with the protein concentration dependence of Trp fluorescence (**Fig. 5a**). Similarly, observation of the unfolded component of Flap+_(I54A)_ is consistent with the lower intensity of Trp fluorescence in this mutant compared to the other mutants (**Fig. 5b**).

Since Flap+_(I54A)_ exhibited NMR spectral feature that contains unfolded fraction, we further characterized how the unfolded fraction changes at 40 µM protein concentration, which is similar to that used for the ITC experiments. The apo form of Flap+_(I54A)_ exhibited a similar spectral feature to that at 3 µM (**Fig. 5e**, black). Addition of pepstatin to the -Flap+_(I54A)_, after recording the apo spectrum, changed the spectral pattern, presumably from that of the apo-dimer form to that of the pepstatin-bound form (**Fig. 5e**, red). Upon further addition of DRV to the Flap+_(I54A)_ solution containing pepstatin, the spectral pattern indicated changes from the pepstatin-bound form to the DRV-bound form (**Fig. 5f**, green). Importantly, the unfolded fraction in the Flap+_(I54A)_, approximately 20%, was not changed upon addition of pepstatin, or further addition of DRV, that mimicked the order of inhibitor competition in the ITC experiments (**Table 2**). Even when DRV was directly added to the folded apo-Flap+_(I54A),_ containing 20% unfolded fraction, the unfolded fraction was neither decreased or increased (**Table 2** and **Fig. S1a**). Only when the protein was folded in the presence of inhibitor did the population of the unfolded fraction became small, 5-8%, indicating that the inhibitors facilitate folding of the protein (**Table 2** and **Fig. S1b**). These NMR resonance patterns of Flap+_(I54A)_ in the presence of DRV are very similar to that of Flap+_(I54)_ which differs by only one residue from Flap+_(I54A)_ (**Fig. S1c**). These observations suggest that the unfolded fraction in the apo-form of Flap+_(I54A)_ is a misfolded component or fragments caused by autoproteolysis (**Fig. S2**), and does not participate in the inhibitor interaction throughout the experiments (**Table 2**).

**Table 2.**
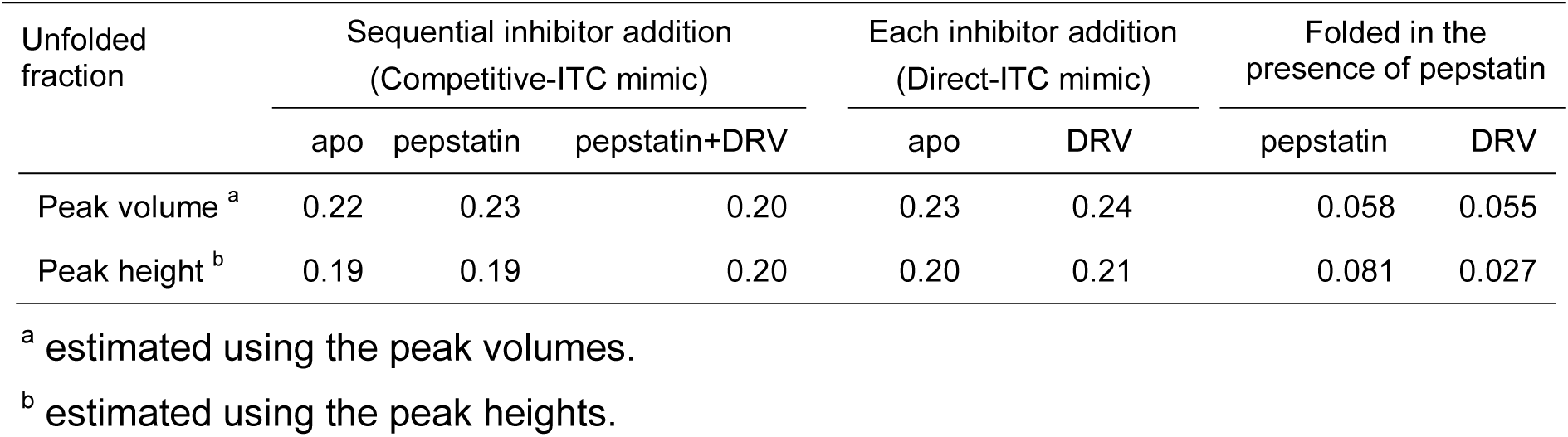
Unfolded fractions in Flap+ (I54A) estimated from Trp indole NMR peak intensity.

## Discussion

In this study, we aimed to understand the molecular mechanism underlying PR-inhibitor interaction, using ITC, NMR and fluorescence spectroscopy. Side-by-side NMR and ITC experiments elucidated the states involved in the thermodynamics changes upon inhibitor-PR interaction. Using this approach, we have made the following overall observations: First, the proteins are folded as a dimer at > µM concentration. Second, Δ*H* of inhibitor interaction with pWT and Flap+_(I54A)_ was more favorable than that of Flap+_(I54)_ and Flap+_(I54V)_, in both DRV and pepstatin interactions. In contrast, *-T*Δ*S* of Flap+_(I54)_ and Flap+_(I54V)_ was more favorable than that of pWT and Flap+_(I54A)_. These observations suggest that evolution from I54V to I54A likely makes the inhibitor-binding thermodynamics of the Flap+ mutant more similar to that of pWT.

Differences in the thermodynamics of the inhibitor interactions with the various PR proteins suggest a previously unrecognized inhibitor-bound intermediate. Specifically, a slow heat release was observed in the pepstatin-DRV competitive ITC experiment of Flap+_(I54A)_ and, less so, of pWT (**Fig. 4a** and **4b**). Importantly, such a slow release was not detected in the direct DRV or pepstatin titration experiments (**Fig. 2a, 2b, 4a** and **4b**). Since the NMR spectra demonstrated that Flap+_(I54A)_ binds to pepstatin, which is subsequently replaced by DRV (**Fig. 5e** and **5f**), the slow process is not due to differences in inhibitor-on/off rates of the proteins or unfolding of the protein, but is due to an intermediate process. If the intermediate is between the apo and inhibitor-bound states, competitive titration would not show such a difference in heat flow, compared to the direct titration, because DRV binds the apo-form in the equilibrium between the apo and pepstatin-bound forms. Thus, we conclude that the intermediate exists but is not located on the PR folding pathway (**Fig. 6a**) and is, instead, located at a step that produces a difference between the direct-titration and competitive titration, i.e, on another inhibitor binding pathway (**Fig. 6b**).

**Figure 6.**
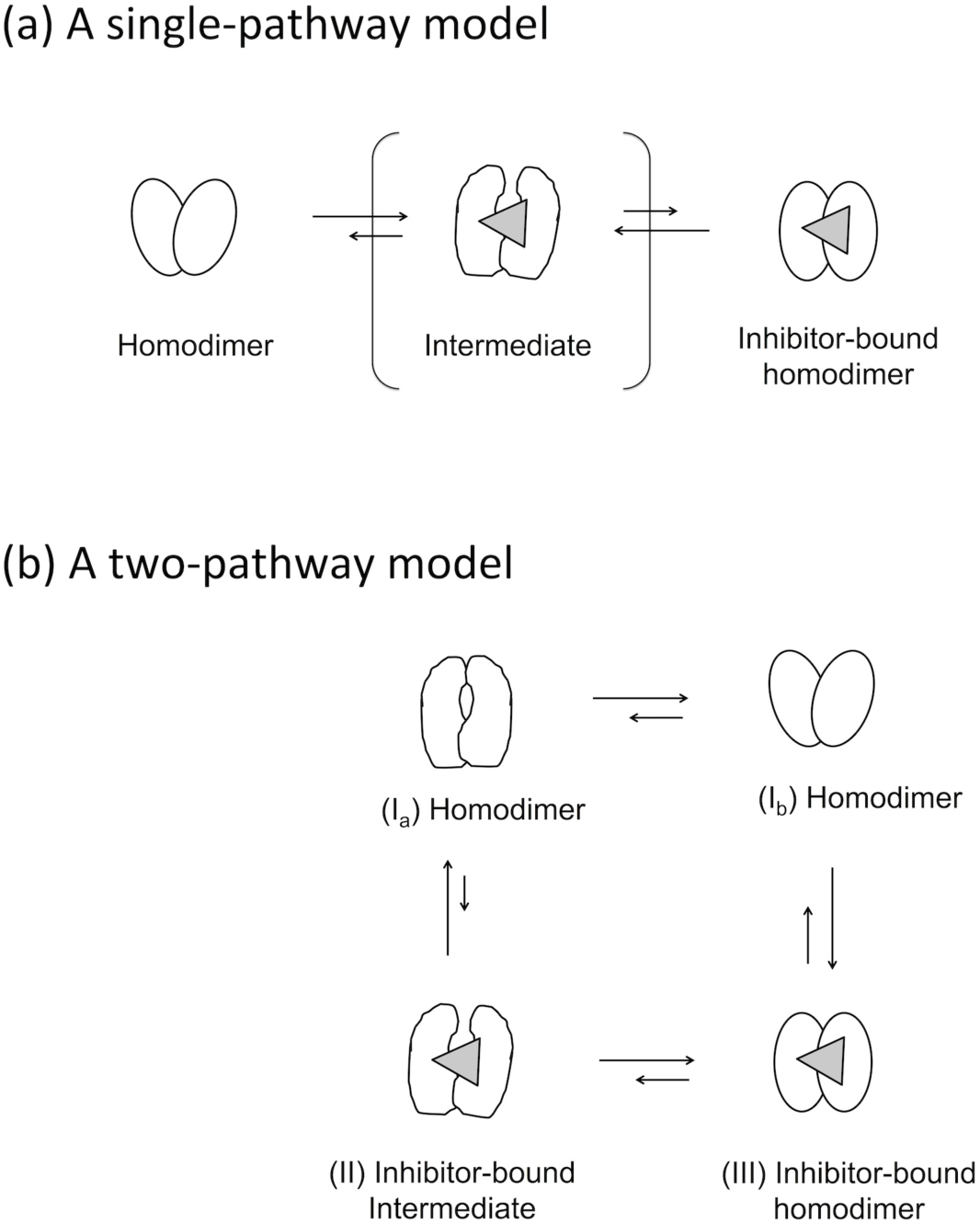
Two models of PR interaction with inhibitors: (a) a single-pathway model and (b) a two-pathway model. Competitive inhibitor titration experiments should result in the same heat flow to that of the direct inhibitor titration in model (a) while competitive inhibitor titration experiments can result in a different heat flow from that of the direct inhibitor titration in model (b).

Based on these observations, we postulate a model that contains an inhibitor-bound intermediate state (state (II) in **Fig. 6b**) in addition to the direct apo-bound pathway (state (I) to (III) in **Fig. 6b**). To place the inhibitor-bound intermediate (II) in the scheme, two states or two different conformers within a state of the apo homodimer, (I_a_) and (I_b_), are assumed (discussed below). The model in **Fig. 6b** explains the heat flow in the competitive titration of Flap+_(I54A)_, and possibly pWT, as follows. In the direct inhibitor-PR titration, PR undergoes either pathway (I)- (III) or (I)-(II)-(III), or both, depending on the mutations. In the competitive ITC experiment, in which PR is filled by a weak binder in state (III), a strong binder binds the low-populated apo form in state (I_b_) and shifts the equilibrium to the strong binder bound form in state (III), in which the direct pathway is (I_b_) – (III). Thus, when (I_b_)-(III) is the major pathway of the direct inhibitor interaction, the competitive experiment will show a heat flow similar to that of the direct interaction. On the other hand, when (I_a_)-(II)-(III) is the major pathway of the direct inhibitor interaction, competitive binding of DRV may occur via the (III)-(II)-(I_a_) pathway (backwards). However, a population that goes through the (III) - (I_b_) pathway (reversed) may also become evident because most of the PR is in the (III) state. In this case, heat flow in the competitive experiments may be different from that of the direct binding experiments, such as observed for Flap+_(I54A)_.

Thermodynamically, we observed that inhibitor interactions of Flap+_(I54A)_ and pWT are Δ*H* driven while inhibitor interactions of Flap+_(I54)_ and Flap+_(I54V)_ are -*T*Δ*S* driven (**Table 1.3**). In the above two-pathway model (**Fig. 6b**), this difference in thermodynamics characteristics may indicate that the inhibitor-bound intermediate path, (I)-(II)-(III), is Δ*H* driven while the direct binding path, (I)-(III), is *-T*Δ*S* driven. In this case, the intermediate path may involve more protein conformational change (Δ*H* driven), while the direct binding path involves burial of hydrophobic inhibitor to the rigid homodimer (-*T*Δ*S* driven).

Structurally, two slightly different dimer forms may exist, as discussed below. Previous studies showed a more open flap conformation in Flap+_(I54V)_ compared to pWT in the apo form [46; 47] and showed that Flap+_(I54V)_ exhibits a slightly different conformation, on average less than 1 Å, from the pWT in the DRV bound form [31; 46; 47] [60]. Thus, the simplest scenario for the two forms in state (I) of our model may be closed- (I_a_) and open- (I_b_) flap forms: when the flaps are more closed, inhibitor binding will need a PR conformational change, which may involve significant enthalpy changes. Another scenario of the two forms in state (I) may instead involve a folding intermediate: the Flap mutations are not simply additive of effects of individual mutation; all four mutations span the direct and indirect dimer interfaces (**Fig. 1**); Flap+_(I54A)_ and pWT showed weaker chemical denaturation compared to Flap+_(I54)_ and Flap+_(I54V)_. The latter model may be valid considering the proposed existence of a PR folding intermediate [48], intermediate-inhibitor bound forms [49], and DRV as a dimer dissociation inhibitor [50; 51]. The two subunits of the PR dimer have multiple direct subunit interaction sites, including the flap region, active-site fireman’s grip (residues 25 to 27), and the N- and C-terminal β-sheet (**Fig. 1**) [35-37]. Indeed, another set of mutations, L10I/ M46I/I54V/V82A/I84V/L90M, that spans both the direct and indirect dimer interfaces, but not a mutant with only the M46I/I54V mutations, also shows the entropy-enthalpy compensation [34]. Although the exact conformation of the intermediate and the binding pathway model is not identified from the current study, our results indicate that an inhibitor-bound intermediate exists in the pWT and Flap+_(I54A)_. Since PR recognizes nine different natural substrates in the Gag-Pol polyproteins [61], it may need flexibility in molecular recognition, i.e., having both a Δ*H* favored intermediate path and a -*T*Δ*S* favored path.

## Conclusion

To understand the mechanism of entropy-enthalpy compensation in drug interaction with HIV-1 PR, we performed ITC experiments and NMR for four PRs at protein concentrations similar to each other. Observed differences in the direct and competitive titration heat flow cannot be explained without assuming an inhibitor-bound intermediate in pWT and Flap+_(I54A)_. Inhibitor interactions of these proteins involve significant conformational changes. Based on our observations, we propose two inhibitor-binding pathways, one without (entropy favorable) and the other with large conformational changes (enthalpy favorable), which may explain entropy-enthalpy compensation detected in the competitive ITC experiments.

## Materials and Methods

### Protease Expression and Purification

HIV-1 PR with the following amino acid sequence, PQITLWKRPL VTIRIGGQLK EALLDTGADD TVIEEMNLPG KWKPKMIGGI GGFIKVRQYD QIPIEIAGHK AIGTVLVGPT PVNIIGRNLL TQIGATLNF, was used in this study. Note, the construct contains four mutations (Q7K, L33I, C67A, C95A) to reduce autoproteolysis and disulfide-bridge formation [57; 58] and L63P polymorphism [31; 62]. This sequence is called pWT in this study, to distinguish it from WT. Flap+_(I54)_ contains mutations L10I/G48V/V82A on the pWT construct (DNA2.0, Newark, CA). Flap+_(I54A)_ and Flap+_(I54V)_ were yielded by introducing a single amino acid mutation, I54A and I54V, respectively, to Flap+_(I54)_. The clones were confirmed by DNA sequencing. We expressed 15N isotope labeled proteins and purified using the protocols published previously [60]. Proteins were folded with 10 mM acetate at pH 6.0, buffer exchanged to a 20 mM sodium phosphate at pH 5.8, and concentrated approximately to 5 or 50 µM (assuming a dimer). As described below, the protein concentration was re-adjusted in individual experiments. Molecular mass of the proteins used for NMR experiments were checked by Bruker QqTOF mass spectrometer. Darunavir was obtained from Celia Schiffer’s group [62]. All protein concentrations in the manuscript are described by assuming the dimer unless otherwise stated.

### Isothermal Titration Calorimetry

Thermodynamics parameters of interaction of pepstatin (Sigma Aldrich, St. Louis, MO) with PR pWT, Flap+_(I54A)_, Flap+_(I54)_, and Flap+_(I54V)_ were determined at 20 °C, by conducting ITC experiments using MicroCal PEAQ-ITC calorimeter (Malvern, Westborough, MA). Buffer of the proteins was exchanged to 20 mM phosphate at pH 5.8 using a pre-equilibrated dialysis cassette (Thermo Fisher Sci., Waltham, MA), with the final protein concentration at 20 – 30 µM, and added 2% DMSO. The dialysis buffer was used to adjust acetyl-pepstatin concentration from 50 mM stock in DMSO solution to 1 mM first, making 2% DMSO condition, and next adjusted 0.3 – 0.6 mM depending on the protein concentration. DRV titration was conducted in a competitive mode, i.e., in protein solution containing the 10-fold pepstatin, 250 – 350 µM DRV prepared from 50 mM stock DRV solution. For comparison purposes, we used pepstatin for all the competitive experiment. For Flap+_(I54A)_, direct DRV titration was also performed to examine the heat flow. In all the experiments, the raw ITC data, after normalizing a constant control heat to zero, and the integrated heat per moles of injected inhibitors, assuming a 1:1 binding model, were plotted. Thermodynamics parameters of PR (dimer) - inhibitor interaction were determined using the Analysis software (Malvern, Westborough, MA).

### Fluorescence spectroscopy

The PR concentration dependence of WT, Flap+_(I54A)_, Flap+_(I54)_ and Flap+_(I54V)_ was examined by recording intrinsic tryptophan fluorescence emission on a FluoroMax-4 spectrofluoremeter (Horiba Scientific, Edison, NJ). Proteins, taken from a -80 °C frozen stock, were diluted, firstly to 4 µM and then step-wise dilutions to record fluorescence emission spectra with an excitation wavelength at 280 nm at room temperature. Emission intensity changes at 350 nm per molar concentration were plotted to compare the structural changes at different protein concentrations. Chemical denaturation of 1 µM pWT, Flap+_(I54A)_, Flap+_(I54)_ and Flap+_(I54V)_ was monitored by recording the Trp emission at 350 nm at different urea (Sigma Aldrich, St. Louis, MO) concentrations in a 20 mM phosphate buffer at pH 5.8 at room temperature. Fluorescence data of the proteins were compared by normalizing the maximum emission at zero urea concentration among all four PRs to 1.0. Note, since the proteases are enzymatically active, each set of experiments was done within 1 hour.

### NMR spectroscopy

All NMR experiments were performed for proteins in 20 mM phosphate buffer at pH 5.8 and 20 °C, recorded either on Bruker Avance spectrometers at either 800 MHz or 900 MHz, and processed by nmrPipe and ccpNMR [63; 64]. ^1^H-^15^N HSQC spectra were recorded for pWT, Flap+_(I54A)_, Flap+_(I54)_ and Flap+_(I54V)_ in the absence and presence of DRV, at 3 µM protein concentration (as a dimer). An additional three sets of ^1^H-^15^N HSQC spectra were recorded for Flap+_(I54A)_: Set 1) three spectra recorded using a single 40 µM Flap+_(I54A)_ sample, in absence of inhibitor, after addition of excess pepstatin, and after adding DRV to the pepstatin-sample, in order to understand the competitive ITC data; Set 2) two spectra recorded using a single 20 µM Flap+_(I54A)_ sample first in the apo form, then after adding excess DRV, to compare with the direct DRV titration experiment; Set 3) two spectra recorded using a single 40 µM Flap+_(I54A)_ protein folded in the presence of an excess amount of pepstatin, and next by adding an excess amount of DRV, to confirm the inhibitor replacement. Inhibitors added for the NMR experiments were 4 – 10 fold excess relative to each protein concentration. Each NMR experiment took 3 – 12 hrs. Fractions of the unfolded component of Flap+_(I54A)_ in experimental sets, (1) - (3), were assessed from Trp indole resonance volume or height.

## ACKNOWLEDGEMENTS

We thank Teresa Brosenitsch for critical reading of the manuscript, Michael Delk for NMR support, Akbar Ali for providing the inhibitors, and Andrew C. Hinck for sharing the ITC instrument of his group for the initial experiments. This study was supported by grants from the National Institutes of Health (P01 GM109767 and S10OD023481).

## Supporting Information

A pdf file that contains the following data is available free of charge: **Figure S1**, NMR spectra overlay of Flap+_(I54A)_ in the absence and presence of inhibitors, and **Figure S2**, mass spectrometry data of the PR and Flap+ mutants.

## Footnotes

1 Abbreviations: pWT, a pseudo wild-type PR containing Q7K, L33I, L63P, C67A, C95A mutations: Flap^+^_(I54V)_, a PR containing L10I, G48V, I54V and V82A mutations on the pWT PR (I54A) backbone; Flap^+^, a PR containing L10I, G48V, I54A and V82A mutations on the pWT PR (I54) backbone; Flap^+^, a PR containing L10I, G48V and V82A mutations on the pWT PR backbone; pepstatin, acetyl-pepstatin; DRV, darunavir.

